# Geometric principles of dendritic integration of excitation and inhibition in cortical neurons

**DOI:** 10.1101/2025.05.16.654465

**Authors:** Soroush Darvish-Ghane, Pankaj Gaur, Graham C.R. Ellis-Davies

## Abstract

We use two-color uncaging of glutamate and *gamma*-aminobutyric acid (GABA) on layer-5 (L5) pyramidal neurons of the cingulate cortex to define how inhibitory control of excitation is controlled by dendritic geometry. Traditionally, GABAergic input was considered as the gatekeeper, thus, receptors closest to the soma were ideally placed to veto excitation. However, recently modeling has advanced several counter-intuitive hypotheses. Since laser uncaging can be directed at will to any position, we used photostimulation to show that inhibition near the sealed end of dendrites distal to excitation is more effective than inhibition near the soma in modulating excitation. Further, dendritic inhibition was found to be branch specific. Finally, we demonstrate that inhibitory input from multiple thin basal dendrites can centripetally elevate to effectively tune distant excitation at the soma. These findings provide direct experimental evidence supporting theoretical predictions based on dendritic cable properties, revealing the critical role of dendritic geometry in shaping the interaction between excitatory and inhibitory neurotransmission.

**Teaser:** By activating excitatory (E) and inhibitory (I) synapses selectively with light we explored the geometric based interaction of E/I transmission in dendrites of pyramidal neurons.

## Introduction

The interaction of excitatory (E) and inhibitory (I) neurotransmission in dendrites is a fundamental component of a neuron’s computational capabilities(*1*). Mapping of postsynaptic GABA (gamma-aminobutyric acid)-ergic receptors locate the expression of these receptors on the dendritic trees, axon initial segment and the soma of pyramidal neurons(*2-4*). From the presynaptic side, different populations of inhibitory neurons target distinct domains of pyramidal neurons(*5-7*). Such structured pre- and post-synaptic inhibitory distribution posits the question of what functional role the spatial distribution of inhibitory receptors has in controlling the output of pyramidal neurons.

Geometrically, dendrites originate from the soma and become narrower as they terminate with a sealed end(*8, 9*). This dendritic sealed end (off-path) acts as a very strong resistor for axial current flow and the increasing diameter of dendrites going towards the soma give the somatic compartment the behavior of a current sink(*8, 9*). In the context of these geometric properties of neurons, depolarization originating from dendrites attenuate sharply from the spines through the dendritic arbors with increasing diameter in the direction to the soma as modeled by the cable equations and later demonstrated by patch clamp experiments (*10-14*). Due to the steep attenuation of excitation, sigmoid dendritic excitatory non-linearities function to amplify signals and operate as coincidence detectors of clustered synaptic activation(*15-19*).

For inhibition, the traditional view is built upon modeling by Wilfred Rall of dendrites as *passive* cables(*9, 11, 20, 21*). A key idea of which is that inhibition is most effective locally and on the path to the soma, thus there is a steep attenuation of the visibility of synaptic inhibition with distance from the synapse(*11, 20, 21*). Therefore, if an inhibitory input conductance change remains highly local, the optimum position for such synapses to act as gatekeepers for neuronal action potential output is to be between the excitatory synapse and the soma (on-path)(*11, 21*). However, there are experimental studies in the literature that are not entirely in agreement with the passive cable models of inhibition(*22-24*). To address these, Gidon and Segev proposed a new measure of “shunt level” to explain the influence of inhibition in terms of neuronal geometry and the positioning of inhibitory synapses relative to the site of excitation(*25*).

Their concept of “shunt level” (SL) describes mathematically how much excitation is required to counter the inhibition, as an indirect measurement of inhibitory strength(*25*). Qualitatively, the SL is modeled to attenuate in dendrites similar to voltage attenuation as described by Rall’s cable equations(*9*). However, the SL attenuation depends on change in the input resistance of the sub-neuronal compartment due to an inhibitory conductance perturbation and the consequential voltage attenuation from the site of an excitatory input to the inhibitory synapse and back in the context of the dendritic sealed end or current sink boundary conditions(*25*). The analytical solutions for SL based on simplified cylinder models considering the off-path boundary condition and the soma as a current sink predicted for an inhibitory synapse on the off-path to be more effective at controlling an excitation that is “on the path” towards the soma(*25*). Additional solutions to the SL predict for inhibition to be visible from a distance to effectively tune distant excitation in neurons(*25*). Further analysis on non-linear behavior of multiple inhibitory conductance perturbations in the context of SL revealed that SL at a neuronal junction innervated by three or more dendritic branches begins to increase non-linearly towards the junction(*25*). However, these results have not yet been experimentally tested in neurons from complex brain tissue.

Testing such ideas in complex neurons from complex brain tissues remains technically challenging. However, uncaging offers the capability of delivering stereotypical inputs of neurotransmitters using either one-photon (1p) or two-photon excitation. A key feature of this technique is that such inputs can be visually designated, and repetitive. In the present study we use two-color uncaging of glutamate and GABA by using violet and green lasers (405 and 514 nm, respectively), two-photon imaging and whole-cell patch-clamp recording in the cingulate cortex brain slices to test the functional effects of E and I receptor positioning inspired by the principles based on solutions to SL. To do this, we synthesized (PEGylated) caged versions of glutamate and GABA and localized E and I synaptic activation on sub dendritic domains using photo-stimulation. We then quantified E/I interaction as a percentage change of the summated E/I potentials(*22*). In the present work, we demonstrate that inhibition at the sealed end (i.e. off-path) of the dendrites is more effective than on-path inhibition in modulating the threshold for dendritic spike (d-spike) generation in a branch specific manner. We also demonstrate that multibranch inhibition of excitation elevates at the somatic junction. Together, these results provide experimental evidence in agreement with the derivation of cable equations(*25*) that implicate the critical role of neuronal architecture in dendritic inhibition.

## Results

### Spatially precise control of dendritic excitation and inhibition with 1P neurotransmitter uncaging

Recently we introduced(*26*) caged glutamate and GABA probes with photosensitive protecting groups based on the laser dye courmain-102 (C102). C102-caged glutamate (C102-Glu) and thio-coumrain-102-caged GABA (SC102-GABA) were photolyzed with exceptional wavelength selectivity using violet and green light(*26*). In the current study, we developed PEGylated version of these compounds to enhance solubility in ACSF (Fig. 1A). The key feature of these new chromophores is that the long wavelength caging chromophore is minimally responsive(*26*) to wavelengths used to photolyze the short wavelength caging chromophore (Fig. 1B). We found these new probes performed as expected when uncaging with visible light on pyramidal neurons in acutely isolated brain slices when compared to probes we have developed previously (Fig. 1B-F). In all experiments we used local delivery of PEG-SC102-GABA (20 *µ*M) or PEG-C102-Glu (500 *µ*M) either near the soma or on isolated dendrites. One-photon (1p) irradiation with a green laser evoked postsynaptic inhibitory currents (1pIPSCs) that increased linearly with laser power (Fig. 1C). Similarly, the irradiation of PEG-C102-Glu with violet laser evoked postsynaptic excitatory currents (1pEPSCs) that increased linearly with laser power (Fig. 1D). In both cases, the slope of dendritic and somatic responses was significantly different (Fig. 1C and D). Next, PEG-C102-Glu was puffed near an isolated dendrite and surveyed for irradiation (0.6 mW) to evoke approximately a 3 mV excitatory postsynaptic potential (1pEPSP). These conditions produced non-linear excitatory sigmoidal events that are characteristic of d(dendritic)-spikes(*10, 23*) (Fig. 1E-F). The lateral resolution of uncaging PEG-C102-Glu resolution the uncaging beam was determined by moving laterally away from the membrane once a d-spike was evoked (Fig. 1G). Normalized 1pEPSPs had a full-width half-maximum (FWHM) value of about 4 *µ*m (Fig. 1G). For the resolution of PEG-SC102-GABA, 1pIPSCs were evoked and the uncaging point was moved laterally away from the dendrite in 5 *µ*m steps (Fig. 1G). The FWHM of the 1pIPSC was about 15 *µ*m (Fig. 1G). Next, we determined the physiological functionality of the compounds. PEG-SC102-GABA irradiation effectively blocked current injection evoked action potentials (APs) and separately, we evoked somatic APs by irradiating PEG-C102-Glu on basal dendrites (Fig. S1, A-B). Next, to determine two-color functionality, we paired green irradiation on the on- or off-path of dendrites with violet (GV) evoked APs initiated from a basal dendrite (Fig. S1, A-C). Green light illumination for 5ms or 0.5ms similarly reduced the probability of APs evoked by dendritic violet irradiation (5ms, on-path: 25% ± 9%, off-path: 21% ± 6%; 0.5ms, on-path: 25% ± 15%, off-path: 27 ± 17%, S1, Fig. S1, C-F).

**Fig. 1.**
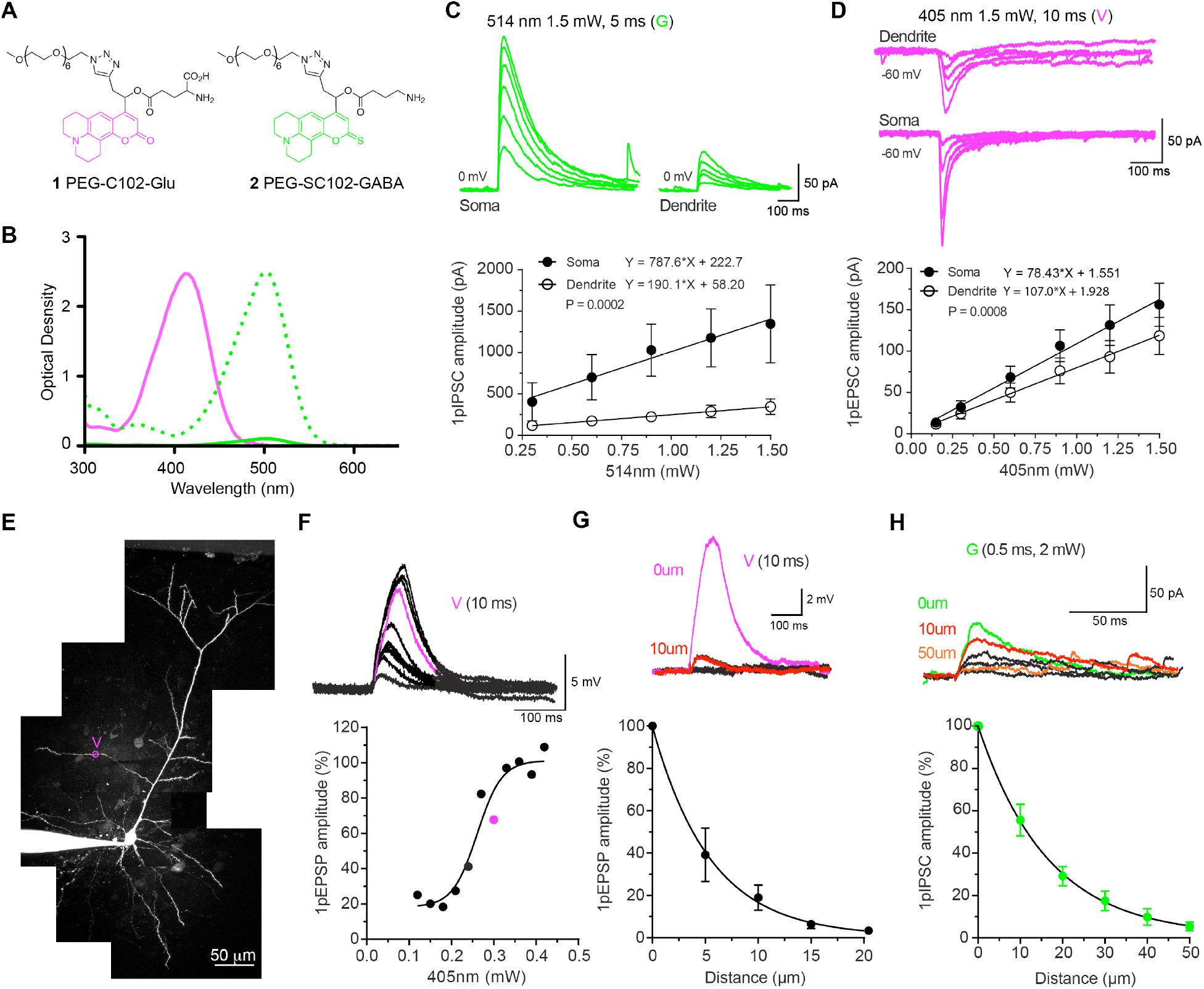
Spatially precise control of dendritic excitation and inhibition with 1P neurotransmitter uncaging. (**A**) Chemical structures of PEG-C102-Glu (**1**) and PEG-SC102-GABA (**2**). (**B**): Absorption spectra of PEG-C102-Glu (violet line) and PEG-SC102-GABA (green solid line) for bath application and a working distance of 2 mm. The green dotted spectrum shows PEG-SC102 normalized to PEG-C102 for comparison. (**C**) Top: representative IPSCs. evoked by irradiating PEG-SC102-GABA with green light (G, 2 wM, 5ms, 514 nm) directed at the soma and isolated dendritic ends. Bottom: Summary plot of 1pIPSC amplitudes vs. power (slope difference; P = 0.0002, n=3/2mice). (**D**) Top: Representative EPSCs evoked by irradiating PEG-C102-Glu with violet light (V, 10 ms, 1.5 mW, 405 nm) directed at the soma and the sealed-end of dendrites in the same neuron. Bottom: current and laser power curves fitted to a linear function. (Slope difference; P = 0.0008, n=4/2mice). (**E**) Maximum intensity projection (MIP) of a two-photon z-stack of a pyramidal neuron filled with Alexa-594 with location of violet irradiation on an oblique dendrite that evoked a non-linear, sigmoid depolarization event (d-spike). (**F**) Top: Sample traces of 1pEPSPs recorded by irradiating PEG-C102-Glu with V (10 ms) on an oblique dendrite of a pyramidal neuron from layer 5 of the ACC. The violet trace depicts supralinear 1pEPSP indicative of a dendritic spike (d-spike). Bottom: The plot of normalized 1pEPSP amplitude against power (mW) fitted with a sigmoid function. (**G**) Top: Sample trace of d-spike and with PEG-C102-Glu irradiation by violet light on the dendritic membrane and subsequent irradiations away from the membrane in 5 *µ*m increments. Bottom: V irradiation resolution as a function of normalized 1pEPSP amplitudes relative to the distance of irradiation beam to the dendrite fitted by a one-phase exponential decay. 1pEPSP reached 50% max value at 5 *µ*m away from the dendrite (n=5/3mice). (**H**) Top: sample traces of 1pIPSCs recorded by PEG-SC102-GABA irradiation by G (0.5 ms, 2 mW) pointed at the dendrite (0 *µ*m) and moved away from the dendrite in 5 *µ*m increments. Bottom: Plot of normalization 1pIPSC amplitudes relative to G distance from dendritic membrane for 0.5 ms. The results were fitted by an exponential decay function.

### Branch specificity of dendritic inhibition

Increases in dendritic diameter at branch points act as current sinks that steeply attenuate voltage going towards the soma and poorly attenuate to a sibling branch(*8*). However, the SL predicts inhibition to sharply attenuate and result in poor spread of inhibition into a thin sibling dendritic branch(*21, 25*). To evaluate branch specificity of inhibition in L5 pyramidal neurons (Fig. 2A), we used violet light to evoke 1pEPSPs on a thin dendritic branch above the branch point and paired it with green light on the same (green), or the sibling (teal) branches and measured the peak amplitude of 1pEPSPs (Fig. 2B). GABAergic uncaging on the same dendritic branch, but not the sibling branch significantly inhibited the amplitudes of dendritic 1pEPSPs (V1: 5.3 mV ± 1.3 mV vs. GV Same: 2.82 mV ± 0.77 mV; P = 0.007, V1 vs. GV Sibling: 4.78 mV ± 1.34 m; P > 0.05, Fig. 2C). We tested branch specificity of inhibition in an alternative manner by the modulation of the d-spike threshold. Dendritic branches (Fig. 2D) were surveyed for regions that evoked d-spikes and GABAergic inputs were paired on the same or the sibling branch (Fig. 2E). Two-color uncaging on the same dendrite significantly increased the threshold for d-spike initiation, while relocation of green irradiation spot to the adjacent sibling branch had no effect on the threshold for d-spike initiation, when compared to the control photolysis with just violet light (V1: 0.37 mW ± 0.05mW vs. GV2 Same: 0.51 mW ± 0.06 mW; P = 0.04, V1 vs. GV3 Sibling: 0.38 mW ± 0.05 mW, Fig. 2F).

**Fig. 2.**
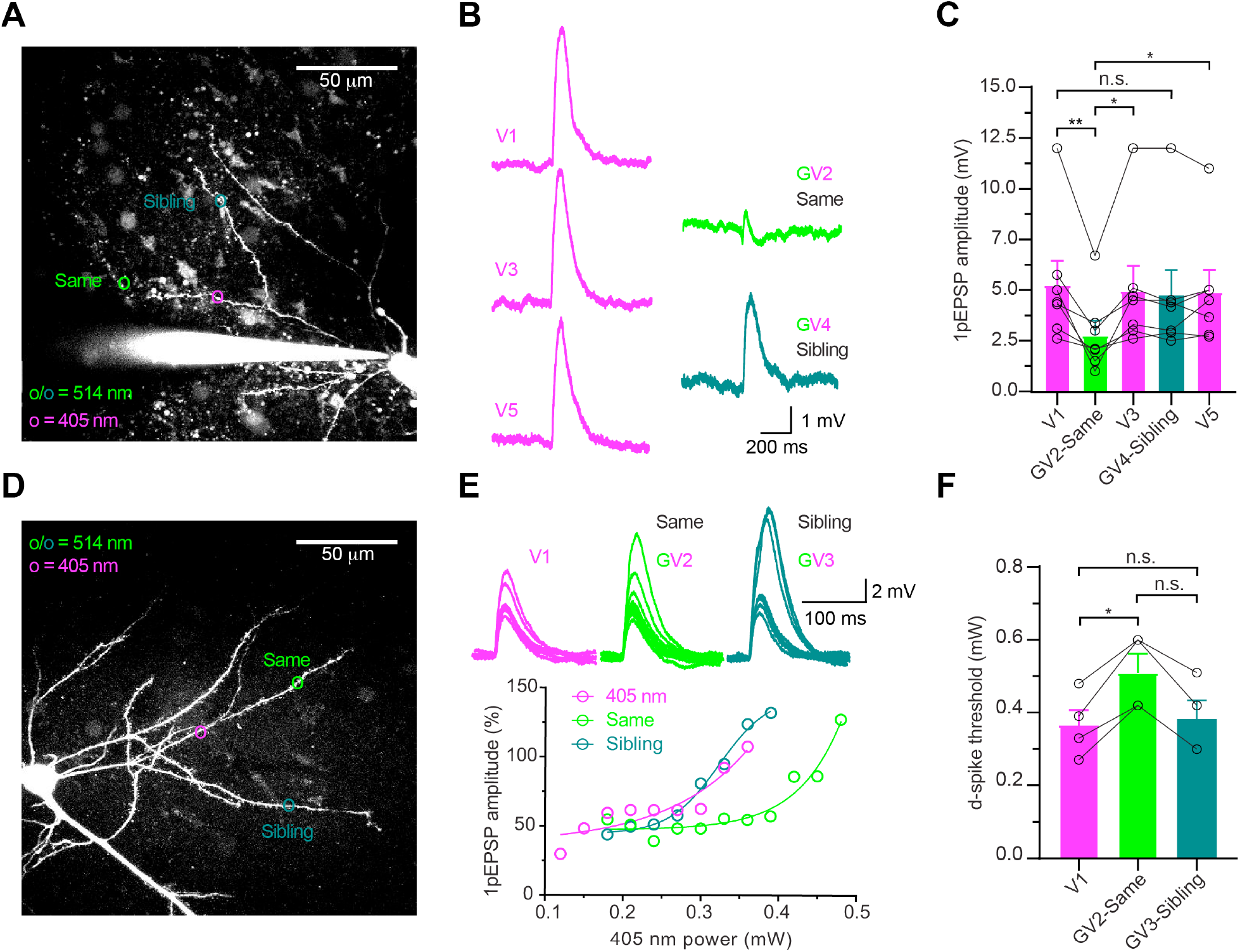
Branch specificity of dendritic inhibition. (**A**) Maximum intensity projection (MIP) of a two-photon z-stack of a pyramidal neuron filled with Alexa-594. Violet (V, 405 nm) and green (G, 514 nm) lasers were used for uncaging. Locations of former (violet circle) and latter (green and teal circles) are shown. A mixture of **1** and **2** were perfused locally. (**B**) The corresponding averaged traces of 1pEPSPs with the V numbering indicating the uncaging sequence. (**C**) Summary of the results for the raw 1pEPSPs in mW comparing inhibition of 1pEPSP when GV experiments were conducted on the same or sibling dendritic branches (Friedman test, P = 0.0034, V1: 5.3 mV ± 1.3 mV vs. GV Same: 2.82 mV ± 0.77 mV; P = 0.007, GV Same vs. V3: 5.0 ± 1.3; P = 0.04, GV Same vs. V5: 4.95 mV ± 1.6 mV; P = 0.017, GV Sibling: 4.78 mV ± 1.34 mV vs. V1; P > 0.05, n=7/6mice). (**D**) MIP of a two-photon z-stack of a pyramidal neuron filled with Alexa 594 and the uncaging locations indicated as **A**. (**E**) The corresponding 1pEPSP traces resulting from a power train with V alone, or GV paired on the same or sibling. (**F**) Summary of the result of the raw power for d-spike threshold when G was paired on the same- or the sibling branch (Friedman test, P = 0.0417, V1: 0.37 mW ± 0.05mW vs. GV2 Same: 0.51 mW ± 0.06 mW; P = 0.04, V1 vs. GV3 Sibling: 0.38 mW ± 0.05 mW; P > 0.05, n=4/4mice). Error bars indicate mean ± SEM.

### More effective off-path dendritic inhibition of d-spike threshold

Having determined the branch specificity of inhibition, we tested whether off-path inhibition is more effective than on-path in modulating d-spikes threshold in basal dendrites. Pyramidal neuron basal dendrites were surveyed for a location that evoked d-spikes with violet illumination (Fig. 3A). Once a d-spike was detected, the violet irradiation was paired with green light illumination, either at on- or off-path locations (Fig. 3B). Both locations significantly increased the threshold for d-spike initiation (Fig. 3C). However, localization of the green laser off-path caused a significantly higher threshold for d-spike initiation compared to the on-path (GV on-path: 0.285 mW ± 0.016 mW vs. GV off-path: 0.339 mW ± 0.025 mW, Fig. 3C). Normalized d-spike threshold power, GV on-path: 124.7% ± 7.7% vs. off-path: 147.7% ± 10.7%, paired t-test, t=2.7, P = 0.0244 (not shown). Furthermore, we tested the threshold with GABA-B antagonist CPG-55845 (3 *µ*M) in the bath and puffing ACSF and observed an increase in threshold when green and violet lights were paired, hence we pooled the results with samples without GABA-B antagonism. Furthermore, analysis of the averaged d-spike amplitudes evoked by GV on-path (12.2 mV ± 0.5mV) marginally decreased in comparison to off-path (12.8 mV ± 0.8mV) and V (13 mV ± 0.6mV) (Fig. 3D). In addition, measured 1pIPSPs from the on- or off-path demonstrated insignificantly different magnitudes of inhibition (on-path: -3.8 mV ± -1.7 mV vs. off-path: -2 mV ± -0.6 mV, Fig. S2). Albeit a lower magnitude of the 1pIPSPS from the off-path may be due to dendritic filtering.

**Fig. 3.**
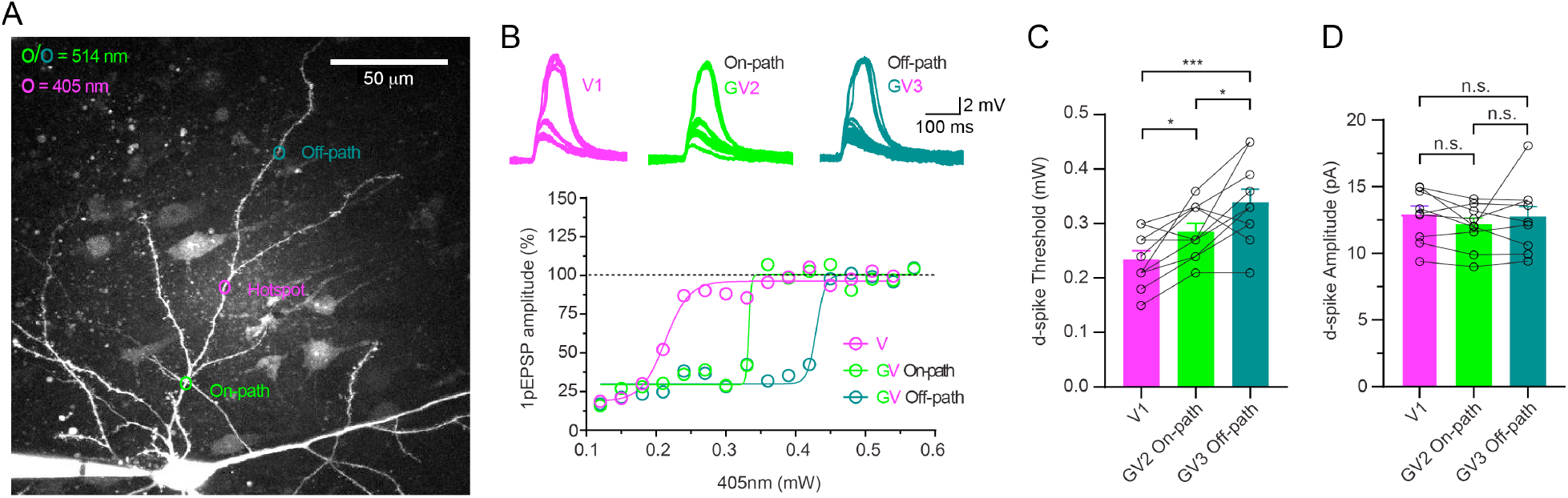
More effective off-path dendritic inhibition of d-spike threshold. (**A**) MIP of a z-stack two-photon image of a pyramidal neuron filled with Alexa-594. Violet (V, 405 nm) and green (G, 514 nm) lasers were used for uncaging. Locations of the former (violet circle) and latter (green and teal circles) are shown. A mixture of **1** and **2** were perfused locally. (**B**) Upper: Representative d-spike voltage traces from a power train at the dendritic hotspot using only V irradiation (violet), and V combined with G irradiation on-path (green) and off-path (teal). Lower: Normalized 1pEPSP amplitudes and laser power (mW) curves fitted into a sigmoid function demonstrate dendritic spike threshold as the steepest part of graph. (**C**) Summary of raw values for the laser power (mW) at the d-spike threshold. G irradiation on the on- or off-path (one-way repeated measure ANOVA, F(1.747, 15.72)=14.24, P = 0.0004, V1: 0.234 mW ± 0.017 mW vs.. GV on-path: 0.285 mW ± 0.016 mW; P = 0.018, V1 vs. GV3 Off-path: 0.339 mW ± 0.025 mW; P = 0.0023, GV on-path vs. GV off-path; P = 0.0353, Holm-Sidak’s multiple comparison, n=10/9 mice). (**D**) Summary of d-spike amplitudes between V and GV trials. (one-way repeated measure ANOVA, F (1.408, 14.08) = 1.378, P = 0.271, on-path: 12.1mV ± 0.47mV vs. V: 13mV ± 0.56mV; p = 0.0598, Tukey’s multiple comparison n=12/10 mice). Error bars indicate mean ± SEM.

### Elevated centripetal inhibition

The SL concept introduced by Gidon and Segev(*25*) appears even more striking when the effects from an analytical solution of how multiple inhibitory conductance from separate branches sum onto their intersection point. Based on their calculations, a minimum of three cylindric branches are required for SL to non-linearly elevate towards a parent junction(*25*). We tested this concept in the following way. First, we evoked 1pEPSPs by violet irradiation on the soma. Then, we paired the same violet stimulation with five points of green irradiation (0.5 ms/point, 2mW) clustered on one basal branch, or distributed on five separate basal branches (Fig. 4A). Our results demonstrate that inhibitory inputs from a thin basal dendrites can shunt depolarization evoked at the soma (Fig. 4A and B). Furthermore, we observed that distributing GABAergic inputs on five branches inhibits somatic excitation significantly more than inhibition clustered on a single basal dendrite (clustered: 4.6 mV ± 0.6 mV vs. clustered: 4.6 mV ± 0.6 mV vs. clustered: 4.6 mV ± 0.6 mV, Fig. 4B). Distributed GABAergic inputs increased the percent inhibition of the averaged and normalized 1pEPSP at the soma in comparison to inhibition clustered on a single dendrite (clustered: G-clustered: 24.8% ± 4.8% vs. G-distributed: 48.34% ± 2.9%, Fig. S3A). In these configurations, distributed multi-branch uncaging evoked significantly higher 1pIPSCs compared to single dendrite clustered inhibition (G1 clustered: 101 pA ± 44 pA vs. G2 distributed: 206 pA ± 62 pA, Fig. 4D). Furthermore, the magnitude of 1pIPSPs between clustered and distributed configurations were not significantly different (clustered: -4 mV ± -2.5 mV vs. distributed: -4.4mV ± -2.11 mV, Fig. S3B and C).

**Fig 4.**
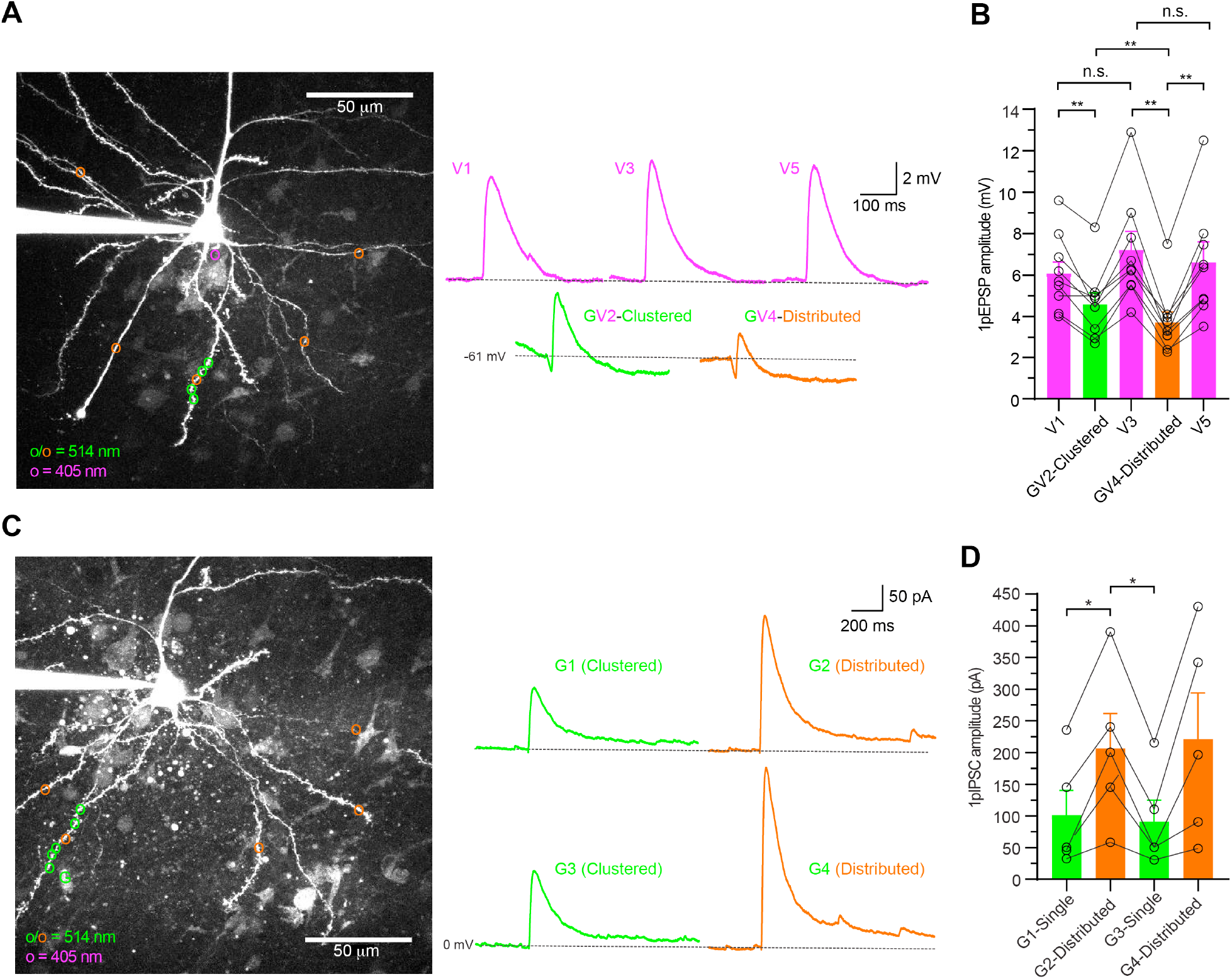
Elevated multi-branch centripetal inhibition. (**A**) Left: MIP of a two-photon z-stack of a pyramidal neuron filled with Alexa 594. Violet (v, 405 nm) and green (G, 514 nm) lasers were used for uncaging. Locations of the former on the soma (violet circle) and latter (clustered, green circles or distributed, orange circles) are shown. Right: the corresponding averaged traces of 1pEPSPs from the depicted neuron with the V numbering indicating the uncaging sequence. A mixture of **1** and **2** were perfused locally. (**B**) Summary of raw 1pEPSPs for V on the soma and GV irradiation in clustered or distributed configuration on basal dendrites. (one-way repeated measure ANOVA, F (1.916, 15.33) = 23.92, P < 0.0001, V1: 6.1 mV ± 0.64 mV vs. clustered: 4.6 mV ± 0.6 mV; P = 0.009, clustered vs.distributed: 3.7 mV ± 0.6 mV; P = 0.005, Tukey’s multiple comparisons test, n=9/7animal. The G irradiations (0.5ms) are separated by 0.012 ms time interval. Sequence of clustered and distributed experiments were altered to account for potential duration dependent effects. See also Fig S3A. (**C**) MIP of a two-photon Z-stack of pyramidal neuron filled with Alexa 594 and the location of five G irradiation (0.5 ms/point) sites clustered on a single dendrite (green circles) or distributed (orange circles),and the corresponding averaged traces of 1pIPSCs from the depicted neuron. Neurons were voltage clamped at 0 mV. (**D**) Summary of raw 1pIPSCs. Distributed GABAergic inputs evoked 1pIPSCs with significantly higher amplitudes compared to inputs clustered on a single dendrite (repeated measure one-way ANOVA, F (1.303, 5.211) = 10.39, P = <0.0001, clustered: 101.4 pA ± 43.58 pA vs. Distributed I: 206.6 pA ± 61.57 pA; P = 0.0389, distributed I vs. Single I: 91 pA ± 61.58 pA; P = 0.0352), n=5/3mice. Error bars indicate mean ± SEM.

## Discussion

The interplay of inhibitory and excitatory synaptic signaling is a topic of long-standing interest in neurophysiology. It was the subject of Sherrington’s Nobel Prize and was studied by Eccles and Katz in the 1950s(*27, 28*), with the concept of an inhibitory shunt being described by the latter in 1953(*28*). Such experimental work was the foundation of mathematical models of the conduction properties of dendrites by theoreticians such as Rall(*9, 29*) and Koch(*11, 21*). The former derived the cable equation for excitatory conduction of dendrites, and the latter extended this model, by suggesting inhibition acts as a “gate keeper” to excitation along the dendritic path to the soma(*11*). In 2012 Segev built on these ideas in a way that started to account for the complex anatomy of inhibitory inputs onto pyramidal neurons(*25*). Importantly, he suggested that inhibitory inputs on distal portions of dendrites could be highly effective in controlling excitation that was more proximal to the soma. However, experimental methods to evaluate these progressive theoretical models fully have remained very challenging.

Chromatically selective uncaging of glutamate and GABA would seem to offer a technical solution to this problem, as synthetic caging chromophore can be tuned chromatically in a facile manner to allow near-zero cross-talk between optical channels(*26, 30, 31*). We used our newly developed caged neurotransmitters (Fig. 1A) to test the hypotheses advanced by derivation of cable equations, using wavelength selective photolysis with violet and green light. This approach is effective because uncaging allows postsynaptic stimulation to be delivered in a stereotypical manner at visually selective spots that are independent of presynaptic release probability.

First, we confirmed that our approach worked in complex brain tissue, in a manner similar to that reported for iontophoresis in cultured neurons (Fig. 2A-C cf.(*32*)). Indeed, we took advantage of using brain slices to extend the findings of Liu(*32*), and showed that the threshold for d-spikes was increased by paired uncaging on the same but not sibling branches (Fig. 2D-E), as predicted by Segev(*25*). Our experimental results confirm the intuitive idea that sibling branches can be electrically isolated, as current must flow along the path of least resistance towards the soma. Such branch specificity of inhibition has been experimentally observed in cultured neurons in the form of EPSP inhibition(*32*) and dendritic calcium transients(*33*) which supports the role of dendritic branches as fundamental units for information processing(*34*). The branch specificity of inhibition precludes the effect of possible GABAergic activation from crowding adjacent dendrites near the soma upon on-path GABA uncaging to tune a local dendritic depolarization.

Further, we explored the non-intuitive hypothesis that inhibitory inputs towards the end of dendrites can modulate excitation very effectively. Indeed, our results reveal that in L5 pyramidal neurons of the ACC, off-path inhibition can be more robust than on-path inhibition in modulating d-spike threshold (Fig. 3) and effectively block somatic action potential generated from dendritic depolarization (Fig. S1). The main reason for such robust off-path modulation arises from the resistance of the dendritic sealed end for axial current flow when compared to the decreasing resistance towards the soma, which acts as a current sink. These ideas and results may go some way to explain why inhibitory synapses have been found on the ends of pyramidal neurons in whole-neuron reconstructions(*4, 5*). By increasing the threshold for d-spike generation, off-path inhibition may be more effective to veto dendrite specific spike generation in physiological conditions. However, on-path inhibition may play a more general role in inhibition of multiple branches converging to the on-path junction.

In contrast, other experimental findings revealed the effectiveness of dendritic inhibition in non-pyramidal granule cells and parvalbumin expressing interneurons of the hippocampus are determined by unique reversal potentials of the on- or off-path dendritic sub-domains that result in either excitation, shunt- or strong hyperpolarizing inhibition, with dendritic geometry having no influence on the effectiveness of inhibition(*35*). Our analysis on the magnitude of 1pIPSPs in different spatial configurations revealed 1pIPSPs that were similar in magnitude between on- or off-path (Fig. S2), and clustered or distributed configurations (Fig. S3). The strong hyperpolarization that was observed (2/12 neurons) were also neuron specific.

The unique demonstration of our technical approach is the non-linear increase in the somatic inhibition from multiple basal dendrite shunt accumulation. The solution to SL predicted non-linear increase of SL towards a junction to maintain a constant SL (see Fig. 4F). However, our results demonstrate that multi-branch inhibition innervating the soma does not stay constant, but rather it elevates at the somatic junction. Intuitively, this effect is based on the increase in total diameter and conduction path of inhibition from multiple branches, resulting in higher current reaching the junction (in our case the soma).

A limitation of this study is that we activate synaptic and extra-synaptic GABA-A receptors. However, we feel that in terms of the modulation of excitation by GABA-A our approach provides an acceptable approximation of synaptic GABA as the kinetics of both synaptic and extra-synaptic are similar when (1p) uncaging is used. To target specific shaft or spine GABARs, and to quantify the precise number of GABARs activated by uncaging, recombinant probes can be used to visualize viable individual synapses in slice and intact brain preparations (*3*).

In our study, we provide unifying experimental results to support a dendrocentric model of inhibition, where neuronal output primarily takes shape in dendritic trees by the non-linear interaction of E/I potentials. Our results, in addition to the SL model provides the blueprint for an optimal location of an inhibitory synapse to be near the end of a dendrite on the same branch as excitation. Recent work on complete reconstruction of excitatory and inhibitory somatosensory pyramidal neurons(*4*) and human dendritic physiology (*36*) have demonstrated that non-linear E/I interaction from multiple branches further endow single neurons with complex computational capabilities. Here, we demonstrate multi-wavelength photochemical control of neurotransmission as a viable method to address questions by testing experimentally counterintuitive dendritic E/I algorithms (*36*).

## Materials and Methods

### Experimental Design

In the present study we sought to activate glutamatergic and GABAergic receptors selectively on dendrite of pyramidal neurons in brain slices. To do so, we designed and synthesized two next-generation caged coumarin based glutamate (PEG-C102-Glu) and GABA (PEG-SC102-GABA) that are activated by green and violet light. These versions of the compounds are made more soluble in artificial cerebrospinal fluid (ACSF) to improve slice penetration and to demonstrate the applicability of the synthesis pathway. With these new compounds, we interrogated principles of dendritic inhibition based on idealized cable properties.

All chemicals were purchased from commercial sources and used as received unless otherwise noted. Reactions were monitored by thin-layer chromatography (TLC) on Merck KGaA glass silica gel plates (60 F254) and were visualized with UV light. NMR spectra were recorded on a Varian 300 MHz or a Bruker 400 MHz NMR spectrometer. The chemical shifts are reported in ppm using the solvent peak as the internal standard. Peaks are reported as: s = singlet, d = doublet, t = triplet, q = quartet, dd = doublet of doublets, m = multiplet. High resolution mass spectral data were obtained using an Agilent G1969A ToF LC-MS (Agilent, Santa Clara, CA, USA).

Reverse-phase chromatography used two systems. Analytical UPLC was carried out with a Waters Acuity Arc (Milford, MA, USA) using a BEH300 C18 column (2 × 50 mm, 1.7 *µ*m particle size) monitored with a 2998 PDA detector. Elution used a linear gradient elution (0-100% acetonitrile, in water with 0.1 % TFA for 6 min). Preparative HPLC was carried out using a Waters PrepLC using an Alltech Altima C-18 column (22 × 250 mm, 5.0 *µ*m particle size) monitored with a 2489 detector at 254 nm. Isocratic elution was used (15 mL min^-1^) with 30% MeCN in water with 0.1% TFA.

### Synthesis

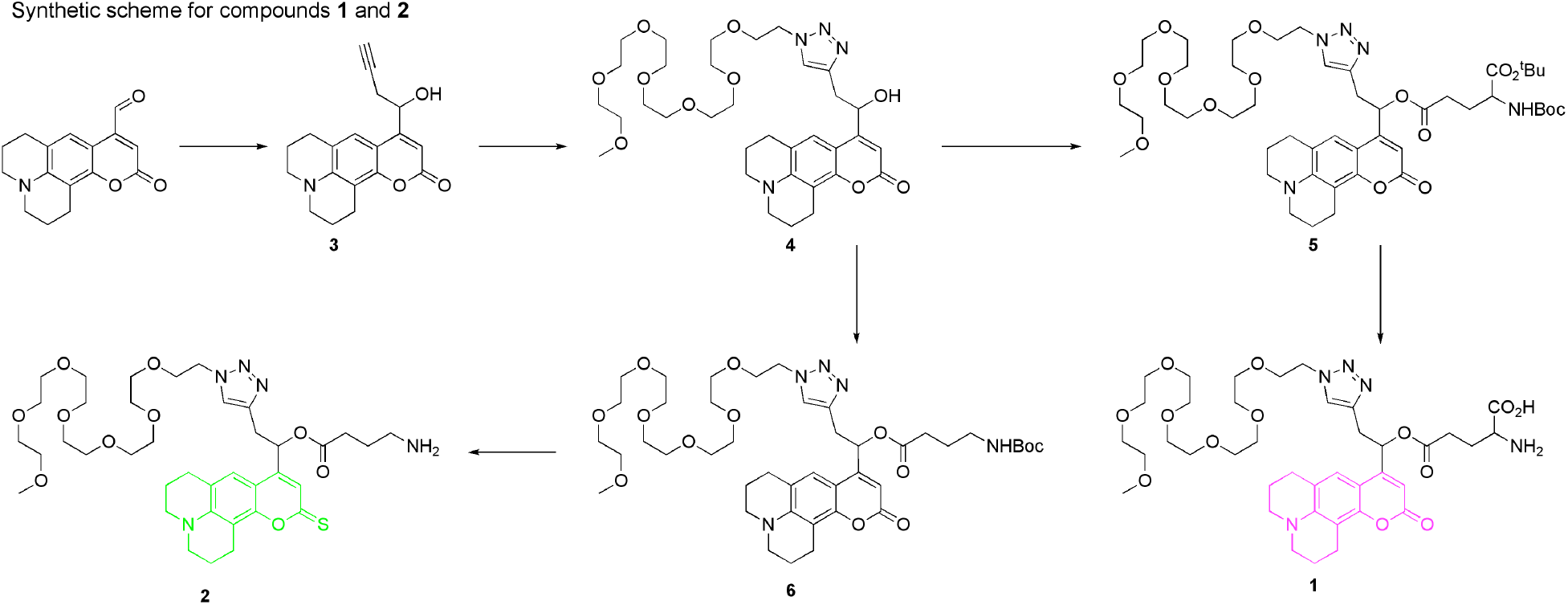

9-(1-hydroxybut-3-yn-1-yl)-2,3,6,7-tetrahydro-1*H*-pyrano[2,3-*f*]pyrido[3,2,1-*ij*]quinolin-11(5*H*)-one (**3**). To a solution of the known C102-aldehyde (7.0 g, 26 mmol) in dry THF (10 mL) was added zinc (8.5 g, 130 mmol) followed by 1,2-diiodoethane (7.3 g, 26.2 mmol). To the reaction mixture NH_4_Cl (75 mL, sat.) was added slowly, followed by propargyl bromide (6.2 g, 52 mmol); then the reaction mixture was heated at 75 °C for 3 h. THF was removed under reduced pressure, and the aqueous phase was extracted with ethyl acetate. The organic phase was washed with water (3x), separated, dried with Na_2_SO_4_, evaporated under reduced pressure and the residue was purified by flash chromatography (40% ethyl acetate in hexanes, v/v) to give 2.50 g (26%) of **3**. ^1^H NMR (CDCl_3_, 300 MHz, δ): 6.97 (s, 1H), 6.25 (s, 1H), 5.13-5.09 (dd, J1 = 6.0 Hz, J2 = 3.0 Hz, 1H), 3.26-3.22 (m, 4H), 2.88-2.82 (m, 2H), 2.79-2.75 (m, 3H), 2.66-2.57 (m, 1H), 2.16 (t, J = 3.0 Hz, 1H), 2.01-1.92 (m, 4H).

^13^C NMR (CDCl3, 75 MHz, δ): 162.86, 155.98, 151.35, 145.71, 120.80, 120.71, 118.23, 107.10, 105.78, 105.04, 79.65, 71.93, 67.84, 67.73, 49.91, 49.46, 27.69, 21.46, 20.53, 20.41. HRMS (m/z): Cal’d for C_19_H_20_NO_3_ [M+H]^+^ = 310.1443, found 310.1443.

9-(2-(1-(2,5,8,11,14,17,20-heptaoxadocosan-22-yl)-1*H*-1,2,3-triazol-4-yl)-1-hydroxyethyl)-2,3,6,7-tetrahydro-1*H*-pyrano[2,3-*f*]pyrido[3,2,1-*ij*]quinolin-11(5*H*)-one **(4)**. To a solution of **3** (0.470 g, 0.152 mmol) and 22-azido-2,5,8,11,14,17,20-heptaoxadocosane (0.492 g, 1.35 mmol) in THF, a freshly prepared aqueous solution of CuSO_4_ (0.024 g, 10 mol%) and sodium ascorbate (0.089 g, 30 mol%) was added, and the reaction mixture was stirred for 45 h. THF was removed and water was removed under reduced pressure and the residue was purified by flash chromatography (2% methanol in dichloromethane, v/v) to give 0.41 g (45%) of **4**. ^1^H NMR (CD_3_CN, 300 MHz, δ): 7.70 (s, 1H), 7.12 (s, 1H), 6.01 (s, 1H), 5.24-5.20 (dd, J = 6, 5 Hz, 1H), 4.48-4.45 (t, J = 6 Hz, 2H), 3.83-3.78 (m, 2H), 3.56-3.52 (m, 22H), 3.48-3.47 (m, 2H), 3.29 (s, 3H), 3.26-3.18 (m, 5H), 3.15-3.13 (m, 1H), 3.02-2.94 (m, 1H), 2.77-2.71 (m, 4H), 1.92-1.88 (m, 4H).

^13^C NMR (CDCl_3_, 75 MHz, δ): 162.89, 157.74, 151.36, 145.36, 143.95, 123.83, 120.97, 118.19, 106.88, 105.98, 104.70, 71.86, 70.52, 59.01, 50.25, 49.89, 49.44, 33.60, 27.75, 21.50, 20.60, 20.48. HRMS (m/z): Calc’d for C_34_H_51_N_4_O_10_ [M+H]^+^ = 675.3605, found 675.3604.

5-(2-(1-(2,5,8,11,14,17,20-heptaoxadocosan-22-yl)-1*H*-1,2,3-triazol-4-yl)-1-(11-oxo-2,3,5,6,7,11-hexahydro-1*H*-pyrano[2,3-*f*]pyrido[3,2,1-*ij*]quinolin-9-yl)ethyl) 1-*tert*-butyl 2-((*tert*-butoxycarbonyl)amino)pentanedioate (**5**). To a solution of **4** (0.110 g, 1eq) in dry DCM EDC (0.192 g, 6.25 eq) and DMAP (0.006 g, 30 mol%) were added, and the mixture was stirred at RT. After 5 min, Boc-L-glutamic acid 1-tert-butyl ester (0.073 g, 1.5 eq) was added and the reaction mixture was stirred at RT for 24h. Reaction mixture was concentrated and the residue was purified by flash chromatography (14:5:1 DCM/EtOAc/MeOH) to give 0.116 g (74%) of **5**.

^1^H NMR (CDCl_3_, 300 MHz, δ): 7.55 (s, 1H), 7.09 (s, 1H), 6.25-6.22 (m, 1H), 5.93 (d, J = 6Hz, 1H), 5.23-5.09 (m, 1H), 4.52-4.47 (m, 2H), 3.85-3.77 (m, 2H), 3.63-3.52 (m, 24H), 3.36-3.31 (m, 4H), 3.27-3.17 (m, 5H), 2.85 (t, J = 6Hz, 2H), 2.78 (t, J = 6Hz, 2H), 2.49-2.39 (m, 2H), 1.99-1.91(m, 4H), 1.50-1.42 (m, 21H) ppm.

^13^C NMR (CDCl_3_, 100 MHz, δ): 173.35, 171.49, 171.21, 170.37, 162.3, 155.47, 153.95, 151.44, 149.32, 146.05, 142.08, 123.53, 120.95, 118.48, 106.98, 105.56, 104.38, 83.31, 82.22, 79.78, 71.92, 70.58, 70.54, 70.5, 70.46, 70.32, 70.25, 69.49, 59.60, 59.02, 53.46, 50.2, 49.95, 49.5, 31.57, 31.48, 31.14, 30.34, 28.32, 27.99, 27.93, 27.92, 27.74, 21.65, 21.46, 20.57, 20.47 ppm.

HRMS (m/z): Cal’d for C_48_H_74_N_5_O_15_ [M+H]^+^ = 960.5181 and obtained = 960.3825.

5-(2-(1-(2,5,8,11,14,17,20-heptaoxadocosan-22-yl)-1*H*-1,2,3-triazol-4-yl)-1-(11-oxo-2,3,5,6,7,11-hexahydro-1*H*-pyrano[2,3-*f*]pyrido[3,2,1-*ij*]quinolin-9-yl)ethoxy)-2-amino-5-oxopentanoic acid

**1**. To a solution of **5** (0.067 g, 1eq) in dry DCM, 2ml TFA was added and the mixture was stirred at RT for 3h. Reaction mixture was concentrated and the residue was purified by reverse phase HPLC (35% ACN/0.1%TFA in H_2_O, Altima C18), then lyophilized to obtain **1** as a yellow powder (43 mg, Yield = 77%).

^1^H NMR (Methanol-d_4_, 300 MHz, δ): 7.87 (s, 1H), 7.22 (s, 1H), 6.33-6.30 (m, 1H), 5.86 (s, 1H), 4.55-4.52 (m, 2H), 4.09-4.04 (m, 1H), 3.83-3.80 (m, 2H), 3.62-3.51 (m, 24H), 3.34-3.28 (m, 9H), 2.82-2.80 (m, 4H), 2.75-2.69 (m, 2H), 2.21-2.14 (m, 2H), 1.99-1.93 (m, 4H) ppm.

^13^C NMR (Methanol-d_4_, 100 MHz, δ): 170.98, 169.96, 163.13, 154.86, 151.28, 146.4, 141.77, 124.5, 120.86, 119.17, 106.35, 105.15, 102.74, 71.45, 70.32, 70.26, 70.09, 70.06, 70.1, 69.98, 69.94, 69.81, 69.79, 69.72, 57.69, 49.94, 49.89, 49.54, 49, 30.76, 30.70, 29.27, 29.22, 27.3, 25.12, 25.06, 21.12, 20.2, 20.06 ppm.

HRMS (m/z): Cal’d for C_39_H_58_N_5_O_13_ [M+H]^+^ = 804.4031 and obtained = 804.4031.

RP-HPLC; retention time = 8.2 min, 35% ACN/0.1%TFA in H_2_O, Flow 20 mL/min, Altima C18, 22×250 mm.

2-(1-(2,5,8,11,14,17,20-heptaoxadocosan-22-yl)-1*H*-1,2,3-triazol-4-yl)-1-(11-oxo-2,3,5,6,7,11-hexahydro-1*H*-pyrano[2,3-*f*]pyrido[3,2,1-*ij*]quinolin-9-yl)ethyl 4-((*tert*-butoxycarbonyl)amino)butanoate **6**. To a solution of **4** (0.105 g, 1eq) in dry DCM, DMAP (0.012g, 60 mol%) and EDC (0.215, 7 eq) were added, and the mixture was stirred at RT for 20 min. Boc-protected GABA (0.05g, 1eq) was added to this mixture and the reaction mixture was allowed to stir at RT for 3h. Reaction mixture was concentrated and the residue was purified by flash chromatography (14:5:1 DCM/EtOAc/MeOH) to give 0.102 g (74%) of **6**.

^1^H NMR (CDCl_3_, 300 MHz, δ): 7.5 (s, 1H), 7.05 (s, 1H), 6.23-6.19 (m, 1H), 5.88 (s, 1H), 4.93 (brs, 1H), 4.46 (t, J = 6Hz, 2H), 3.8-3.73 (m, 2H), 3.6-3.55 (m, 22H), 3.51-3.47 (m, 2H), 3.32-3.28 (m, 4H), 3.24-3.17 (m, 5H), 3.08-3.04 (m, 2H), 2.81 (t, J = 6Hz, 2H), 2.74 (t, J = 6Hz, 2H), 2.36 (t, J = 6Hz, 2H), 1.94-1.88 (m, 4H), 1.75-1.7 (m, 2H), 1.38 (m, 9H) ppm.

^13^C NMR (CDCl_3_, 100 MHz, δ): 171.82, 162.31, 155.99, 153.97, 151.40, 146.03, 142.10, 123.51, 120.97, 120.76, 118.44, 106.91, 105.40, 104.25, 79.03, 71.86, 70.61, 70.53, 70.5, 70.49, 70.44, 70.39, 69.44, 59.09, 58.88, 50.16, 49.44, 31.40, 28.47, 28.30, 27.71, 25.07, 21.40, 20.44 ppm.

HRMS (m/z): Calculated for C_43_H_66_N_5_O_13_ [M+H]^+^ = 860.4657 and obtained = 860.3440.

2-(1-(2,5,8,11,14,17,20-heptaoxadocosan-22-yl)-1*H*-1,2,3-triazol-4-yl)-1-(11-thioxo-2,3,5,6,7,11-hexahydro-1*H*-pyrano[2,3-*f*]pyrido[3,2,1-*ij*]quinolin-9-yl)ethyl 4-aminobutanoate **2**. To a solution of **6** (0.014 g, 1eq) in dry toluene, Lawesson’s reagent (0.022g, 3.5eq) was added and the mixture was heated at reflux temperature for 18 h, under Argon atmosphere. The solvent was removed under high vacuum, and the crude material dissolved in dry 3 mL DCM, followed by addition of TFA (5% v/v) and the mixture was stirred under Argon at RT for 1.5 h. DCM was removed under reduced pressure and TFA was removed under high vacuum. The residue was purified by reverse phase HPLC (40% ACN/0.1%TFA in H_2_O, Altima C18), and then lyophilized to obtain **2** as an orange compound (4 mg, Yield = 32%).

^1^H NMR (Methanol-d_4_, 300 MHz, δ): 7.89 (s, 1H), 7.34 (s, 1H), 6.78 (s, 1H), 6.36-6.32 (m, 1H), 4.58-4.55 (m, 2H), 3.89-3.84 (m, 2H), 3.64-3.60 (m, 22H), 3.56-3.52 (m, 2H), 3.39-3.34 (m, 9H), 2.95-2.83 (m, 6H), 2.61-2.53 (m, 2H), 2.04-1.85 (m, 6H) ppm.

^13^C NMR (Methanol-d_4_, 100 MHz, δ): 196.17,171.22, 154.63, 146.90, 146.77, 141.93, 124.36, 121.28, 120.74, 117.26, 107.46, 105.74, 71.38, 70.12, 70.03, 69.96, 69.88, 69.85, 69.80, 69.72, 69.63, 69.51, 68.93, 57.69, 49.73, 49.62, 49.20, 38.66, 30.81, 30.28, 27.44, 22.27, 20.89, 20.08, 19.95 ppm.

HRMS (m/z): Cal’d for C_38_H_58_N_5_O_10_S [M+H]^+^ = 776.3904 and obtained = 776.3905. RP-HPLC; retention time = 19.44 min, 40% ACN/0.1%TFA in H_2_O, Flow 10 mL/min, Altima C18, 22×250 mm.

### Aqueous stability

Aqueous stability of PEG-C102-Glutamate. **1** was dissolved in 40 mM HEPES with 100 mM KCl (pH = 7.24) and allowed to stand at RT. Hydrolysis was monitored by UPLC. After 4 days, 56% PEG-SC102-GABA remained, corresponding to a t_1/2_ of about 4d.

Aqueous stability of PEG-SC102-GABA. **2** was dissolved in 40 mM HEPES with 100 mM KCl (pH = 7.24) and allowed to stand at RT. Hydrolysis was monitored by UPLC. After 9 days, 37% PEG-SC102-GABA remained, corresponding to a t_1/2_ of about 8d.

### Animals, slice preparation and electrophysiology

All animal procedures were carried out in accordance with NIH and Mount Sinai IACUC care guidelines. C57BL/6 male and female mice were used in equal numbers. Mice (between 5 and 6 weeks of age) were used for all experiments. Mice were kept on a 12-hour light/dark cycle in conventional housing and access to food and water.

### Acute slice preparation

Coronal brain slices (300 *µ*m) from the anterior cingulate cortex were prepared from 5-6 weeks old C57/BL6 mice of both sexes from Jackson Laboratories (Bar Harbor, ME, USA). Animals were anesthetized with isoflurane and decapitated. Slicing was performed with a vibratome (Leica VT1200) in ice-cold oxygenated (95% O_2_; 5% CO_2_) artificial cerebrospinal fluid (ACSF) containing (in mM): NaCl 120, KCl 3, NaHCO_3_ 25, NaH_2_PO_4_ 1.25, CaCl_2_ 1.2, MgCl_2_ 1.2, glucose 10, sodium pyruvate 3, and ascorbic acid 1. Subsequently slices were stored in ACSF at room temperature for at least one hour. All recordings were performed at room temperature.

### Electrophysiological recordings

An Olympus BX-61 microscope was used to visualize pyramidal cells with IR differential interference contrast microscopy. Patch-clamp recordings were performed from morphologically and electrophysiologically identified L5 pyramidal neurons of the ACC. Voltage- and current-clamp recordings were performed with a EPC10 amplifier and Patchmaster software (Heka Instruments, Bellmore, NY, USA). Current and voltage signals were filtered at 10 kHz and digitized at 20 kHz. Patch pipettes were prepared with thin-wall glass (1.5 O.D., 1.1 I.D.). Pipettes had resistances ranging from 4 to 6 MΩ and the capacitance was fully neutralized prior to break in. For 1pEPSP recordings, the standard potassium based intracellular solution contained (in mM): 145 K-gluconate, 5 NaCl, 1 MgCl_2_, 0.2 EGTA, 10 HEPES, 2 Mg-ATP, and 0.1 Na_3_-GTP (adjusted to pH 7.2 with KOH). For 1pIPSC (voltage-clamped at 0 mV) and EPSC (voltage-clamped at -60 mV) recordings, a cesium methanesulfonate based internal solution was used: (in mM) 120 Cs-MeSO_3_, 5 NaCl, 1 MgCl_2_, 0.5 EGTA, 2 Mg-ATP, 0.1 Na_3_GTP, 10 HEPES, and 5 QX314 (pH adjusted to 7.3 with CsOH, 290 mOsm). For two-photon imaging, Alexa 594 (40 *µ*m) was loaded via the patch pipette. Once the whole cell configuration was obtained, at least 20 minutes was allowed for Alexa 594 to diffuse into the dendritic arbors. Electrophysiology recording traces of graded potentials in the figures are average of a minimum of four consecutive traces.

### Two-photon imaging and two-color, one-photon glutamate and GABA uncaging

An Olympus BX-61 microscope (Penn Valley, PA, USA) fitted with a Prairie Technologies (Middleton, WI, USA) Ultima dual-galvo scan head using Prairie View (PV) 5.7. Imaging used a Coherent Ultra-II laser at 1060 nm through a water-immersion lens (60X, 1.0 NA; Olympus, LUMPLFLN 60X). Laser beam intensity was controlled with electro-optical modulator (model 350–50; Conoptics). Uncaging was effected using a Coherent Galaxy 8 (136384), controlled with an Obis laser box (1228877), that combined pig-tailed Obis 532-nm (1276599) and 405-nm (1236438) lasers. The combined output from the Galaxy 8 was sent to the second Ultima scan head via a single-mode FP optical patch cable via a Prairie Technologies laser launch. Data acquisition and timing of uncaging was controlled with PV 5.7. The stocks of the caged neurotransmitters was stored in at -20 °C in acetonitrile. For sample preparation, the compounds were air dried and diluted in puffing HEPES ACSF (in mM): 125 NaCl, 2.5 KCL, 10 HEPES, 2 CaCl_2_, 1 MgCl_2_, 1.25 NaH_2_PO_4_, 25 Glucose, 1 Na Ascorbate (pH adjusted to 7.4 with NaOH). The compounds dissolved in puffing ACSF then loaded into a pipette with tip resistance of 3 MΩ. Once a region of interest for uncaging was determined, the puffing pipettes were positioned and puffed in 500 ms pulses, followed by 10 s of wait time for diffusion for each uncaging trial. Imaging was used a 512×512 grid at 4 ms per pixel dwell time. The z spacing was typically 0.7 mm. Maximum intensity projections (MIP) of z-stacks were made in ImageJ.

### Quantification and statistical analysis

Electrophysiology recordings were analyzed in Fitmaster software (Heka Instruments, Bellmore, NY, USA). Statistical analysis was performed in GraphPad Prism. For fitting analysis, standard one-phase decay and sigmoid (variable slope) functions provided by GraphPad Prism were used. Statistical comparisons were performed on raw values or normalized means as stated in the manuscript. Shapiro-Wilk normality tests were performed to determine the appropriate parametric or non-parametric tests, and the appropriate multiple comparison post hoc tests were carried out for comparison of three or more groups. Statistical details and the number of animals used can be found in figure legends. Unless otherwise stated, results are presented as mean ± SEM.

## Acknowledgments

We thank Profs. Lu-Yang Wang, Deanna Benson and Masanori Matsuzaki for their comments on earlier version of the manuscript.

## Funding

This work was supported by grants from the NIH (NS111600) and HFSP to GCRED.

## Author contributions

Conceptualization: GCRED, SDG Methodology: SDG, PG, GCRED. Investigation: SDG, PG. Visualization: SDG, GCRED. Supervision: GCRED. Writing original draft: SDG, GCRED. Writing—review & editing: SDG, GCRED, PG.

## Competing interests

Authors declare that they have no competing interests.

## Data and materials availability

All data are available in the main text or the supplementary materials. The compounds, generated materials and data sets during the current study are available from the corresponding author upon request. The study did not generate any original code.

## Notes

### Competing Interest Statement

The authors have declared no competing interest.

