## Supplementary material for "Geometric principles of dendritic integration of excitation and inhibition in cortical neurons": SI

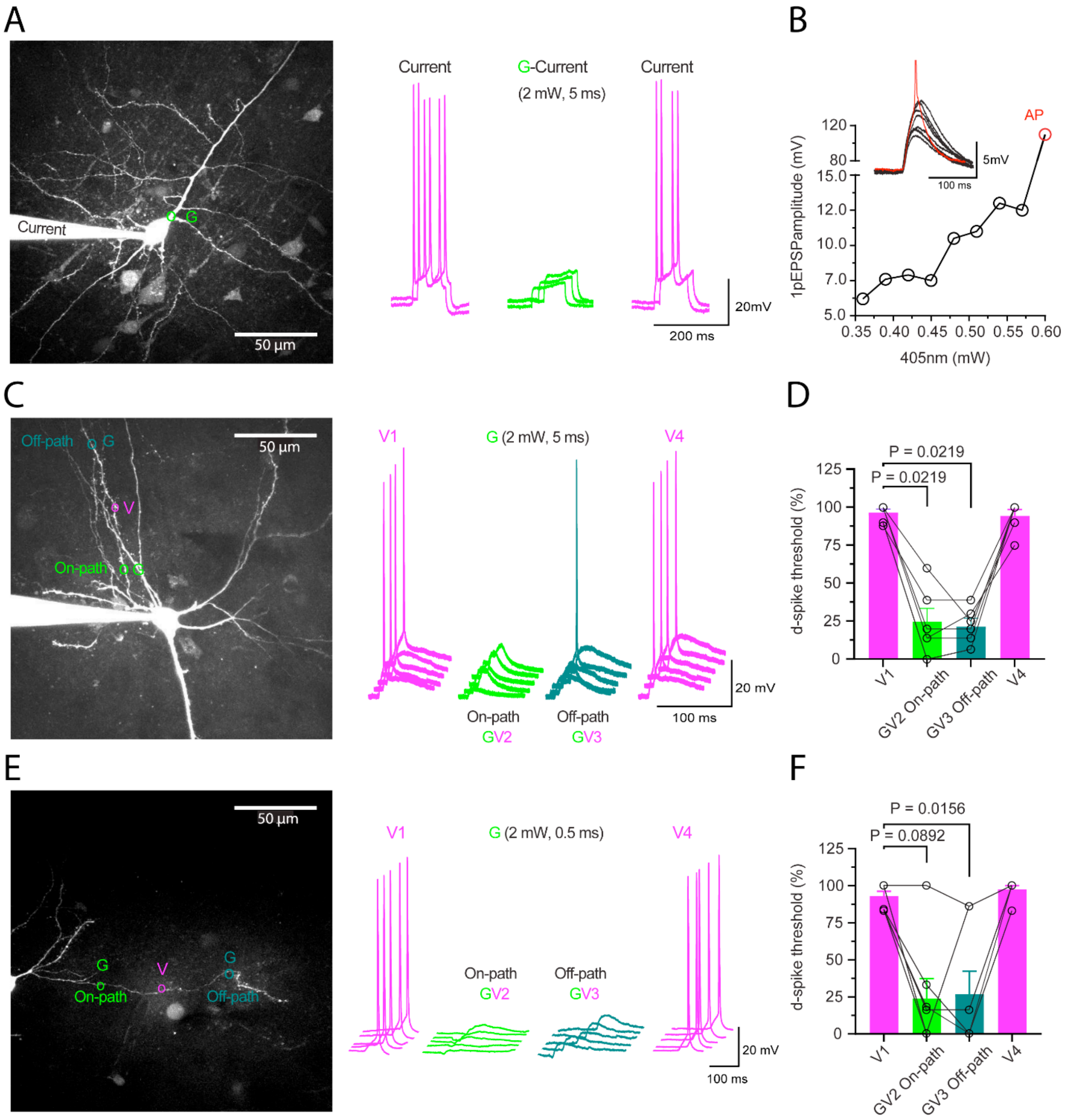


**Fig. S1. Two-color control of dendritic excitation and inhibition in somatic spiking, related to Fig 1.**

(**A**) Maximum intensity projection (MIP) of a two-photon z-stack two-photon image of a pyramidal neuron filled with Alexa 594 with the location of G irradiation on the main somato-dendritic branch and the corresponding sample trace of somatic action potential induced by current injection (200 pA) and G irradiation of PEG-SC102-GABA (green) to block somatic firing.

(**B**) Sample traces of 1pEPSPs and somatic action potential evoked by V irradiation of PEG-C102-Glu on basal dendrite of a layer five pyramidal neuron from the anterior cingulate cortex. Increasing V power (mW) induced a single action potential (red trace).

(**C**) Maximum intensity projection (MIP) of a two-photon z-stack image of a sample neuron depicting locations of V irradiation and G irradiation uncaging on the on- or the off-path dendritic loci. Middle: The corresponding sample traces of somatic action potentials (violet) evoked by V irradiation (10 ms) on basal dendrite and vetoing of the somatic spike with paired G irradiation on the on- or off-path dendritic loci. V duration was 10ms and power level were titrated to evoke single action potential by V irradiation of PEG-C102-Glu on a basal dendrite.

(**D**) Summary of the probability of action potential firing with V irradiation alone or paired with G (5ms). Paired GV irradiation 10ms apart on both the on- or off-path significantly reduced somatic AP probability (Friedman test, P =0.0001, V1: 96% ± 3% vs. GV on-path: 25% ± 10%; P = 0.0219, V1 vs. off-path: 21% ± 6%; P= 0.0219, V1 vs. V4; P>0.05). GV and V4 recovery trials were compared to V1 using Dunnett's case comparison.

(**E**) Maximum intensity projection (MIP) of a two-photon z-stack of a sample neuron depicting location of V and G (0.5 ms) for spatially defined two-color control of dendritic excitation and inhibition. Corresponding sample traces of somatic action potentials (violet) evoked by V irradiation on a basal dendrite and vetoing of the action potential when G was paired with V.

(**F**) Summary of action potential probability in response to V irradiation or GV paired irradiation (0.5 ms, 2 mW) with G on the perisomatic (on-path) or dendritic (off-path) locations. (Friedman test, p = 0.0005, on-path: 24% ± 16% vs. off-path: 27% ± 19%, p >0.05, n=6/5mice). GV and V4 trials were compared to V1 (baseline) using Dunnett's case comparison. Error bars indicate mean ± SEM

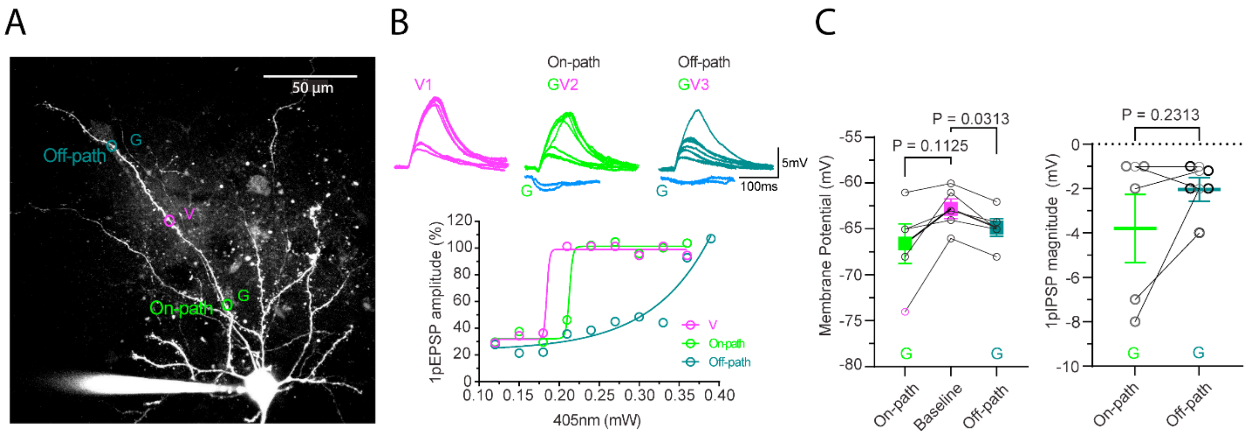


**Fig. S2. One-color 1p GABA uncaging on the dendritic on- or off-path loci, related to Fig 3.**

(**A**) Maximum intensity projection (MIP) of a two-photon z-stack image of a pyramidal neuron filled with Alexa 594 and the locations of V and G (on- or vs-path) irradiation on a basal dendrite.

(**B**) The corresponding traces of 1pEPSPs and 1pIPSP (blue) at the respective On- or off-paths. Bottom: the function of normalized 1pEPSP against 405 nm beam power (mW) fitted with a sigmoid function. Blue traces are 1pIPSP evoked by G irradiation on the respective dendritic locations.

(**C**) Summary of resting membrane potentials at baseline and at lowest value, 20ms~ post G irradiation at the respective on- or off-path loci. Left: No significant difference between IPSPs evoked by G irradiation on the on- or off-path dendritic loci (repeated measure one-way ANOVA, F (1.148, 4.593) = 5.192, P = 0.07, baseline: -62.8 mW ± -1.2 mV vs. on-path: -64.8 mV ± -1.0 mV; P = 0.03, n=6/6 mice). Right: The magnitude of 1pIPSPs GV trials were compared to resting membrane potential with Dunnett's multiple comparison. Error bars indicate mean ± SEM
